# In-depth investigation of genome to refine QTL positions for spontaneous sex-reversal in XX rainbow trout

**DOI:** 10.1101/2024.10.26.620424

**Authors:** Audrey Dehaullon, Clémence Fraslin, Anastasia Bestin, Charles Poncet, Yann Guiguen, Edwige Quillet, Florence Phocas

## Abstract

Sex determination is a flexible process in fish, controlled by genetics or environmental factors or a combination of both depending on the species. Revealing the underlying molecular mechanisms may have important implications for research on reproductive development in vertebrates, as well as sex-ratio control and selective breeding in fish. Phenotypic sex in rainbow trout is primarily controlled by a XX/XY male heterogametic sex determination system. Unexpectedly in genetically XX all-female farmed populations, a small proportion of males or intersex individuals are regularly observed. This spontaneous masculinisation is a highly heritable trait, controlled by minor sex-modifier genes that remain unknown, although several QTL regions were detected in previous studies. In this work we used genome-based approaches and various statistical methods to investigate these QTL regions. We validated in six different French farmed populations DNA markers we had previously identified in a different commercial population on chromosomes Omy1, Omy12 and Omy20. We also identified functional candidate genes located that may be involved in spontaneous masculinisation by reducing germ cell proliferation and repressing oogenesis of XX-rainbow trout in the absence of the master sex determining gene. In particular, *syndig1*, *tlx1* and *hells* on Omy1, as well as *khdrbs2* and *csmd1* on Omy20 deserve further investigation as potential sex-modifier genes to precise their functional roles as well as their interaction with rearing temperature. Those findings could be used to produce all-female populations that are preferred by farmers due to a delayed maturation of females and higher susceptibility of male trout to diseases.

## Introduction

Sex reversal has been defined as a mismatch between the phenotypic and the genetic sex of an individual [1]. In fish, sex reversal can be induced by hormonal treatments (estrogens to feminize and androgens to masculinize) during critical fry stages [2, 3]. Manipulating the sexual phenotype of fish has been used to prevent early maturity and reproduction, as well as increase production in various cultured species or create subsequent sterile generations for fisheries management [3]. Sex-reversed individuals can be used to produce populations that are genetically all-male or all-female (also known as ‘monosex’). Rainbow trout (*Oncorhynchus mykiss*) is a worldwide cultured salmonid species whose production is frequently based on all-female stocks. The production of large rainbow trout over 1 kg body weight has become a growing commercial interest with the development of a market for fresh and smoked fillets. A major bottleneck for this commercial market is the early maturation of males (1-2 year) relative to females (2-3 years) that causes stopped growth, increased sensitivity to pathogens (*Saproleignia* spp.), and reduced flesh quality. Therefore, all-female rainbow trout stocks are often preferred in aquaculture. These stocks are currently produced by crossing hormonal sex-reversed genetically XX-females, known as neomales (XX males), with XX-females. The growing concerns or potential risks to human and environmental health make the prevalent approach of producing neomales via the oral application of the hormone 17-alpha methyltestosterone less sustainable. The finding of a sex control alternative to hormonal treatment based on a safe, consumer and environmentally friendly method is a major challenge for the production of all-female populations [4].

Sex determination and sex differentiation mechanisms in fish are diverse and complex. Understanding sex-differentiation processes requires to unravel the origin and developmental pathways of cells and organs involved in the formation of the primordial gonad. During primary sex determination in vertebrates, the primordial bipotential gonad commits to either an ovary or a testis developmental fate. The structure of fish gonads is similar to that of other vertebrates, with germ cells and associated supporting somatic cells intermixed. Within both ovary and testis, a clear distinction between germ and somatic cells can be made, with the former having the potential to mitotically divide and enter meiosis, and the latter differentiating into associated structural and endocrine cell types. In teleosts, primordial germ cells (PGCs) are established and specialized at the early blastula stage and migrate to the genital ridge during embryonic development [5].

Most of the teleosts are gonochorists meaning that all individuals within a species develop either as males or females and remain the same throughout their lives. Various systems of sex determination have been observed in gonochoristic fishes [1], spanning from a genetic sex determination system (GSD) with a genetic monofactorial system, XX/XY or ZZ/ZW, to different multifactorial systems, or environmental sex determination (ESD). During sex differentiation, the testis and ovary pathways are mutually antagonistic and compete for control of gonad fate. Sex reversal occurs when the sexual trajectory of the gonad is changed to the opposing pathway, switching the sexual phenotype of the organism to the opposite sex. The change in trajectory often occurs because of a failure to maintain the initiated pathway or a failure to repress the opposite pathway [6]. Once the gonad has committed to a particular fate, it begins to produce sex-specific hormones that will drive the secondary process of sex differentiation in which the somatic tissues of the organism differentiate toward one sex or the other. The developmental pathway leading to steroid production in gonadal somatic cells requires complex regulation of multiple genes [7] involved in the differentiation of steroid-producing cells. Sex steroids have local and direct effects on germ-cell development, but also act as endocrine hormones to influence other cell types and organs involved in sex differentiation.

Over the last two decades evidence has accumulated that both somatic and germ cells play critical roles in fish gonadal differentiation. However, the exact mechanisms and factors involved remain to be elucidated [8, 9]. As evidenced from studies in mammals [10], somatic cells may first differentiate in response to the activation of master sex-determining genes, and subsequent differentiation of PGCs into male or female gametes follow in the differentiating testis and ovary in response to signals derived from surrounding somatic cells. Alternatively, it is possible that, first, PGCs interpret internal genetic or external environmental cues and directly transform into spermatogonia or oogonia; then, the surrounding somatic cells may be induced by the germ cells to differentiate accordingly to provide an appropriate hormonal environment for further gonadal differentiation [11]. It is thought that control of estradiol synthesis could play a key role not only for ovarian, but also for testicular differentiation and sex change in fish [12]. This working hypothesis states that the gonadal aromatase gene, *cyp19a1a*, up-regulation would be needed not only for triggering, but also for maintaining ovarian differentiation; and that *cyp19a1a* down-regulation would be the only necessary step for inducing a testicular differentiation pathway. In Atlantic salmon as well as in loach and goldfish, it was shown that germ cells are not essential for gonadal sex differentiation [13]. Germ cell-free female gonads develop an ovarian somatic structure in these species, unlike in zebrafish or medaka where all germ cell-free fish develop somatic testes.

Evidence from both the zebrafish and medaka models [14, 15, 16, 17] suggests important feminizing roles for the germ cells in gonadal differentiation and thus corroborates this alternative hypothesis. In particular, when germ cells are ablated in medaka, XX fish show female-to-male sex reversal, while XY fish exhibit male-to-female sex reversal [16]. Any disruption of this germ cell/somatic cell cross-talk would then disrupt ovarian somatic cell differentiation, leading to an absence of estrogen synthesis and a subsequent masculinization. More subtle regulations not involving the complete loss of germ cells, but instead some differential germ cell proliferation rates, could be also suggested as important triggers of gonadal sex differentiation. In this respect, it is worth to note that the Japanese medaka sex determining gene, *dmrt1Y*, has been found to be an inhibitor of germ cell proliferation [18] and is only expressed during testicular differentiation [19].

In species with genetic sex determination (GSD), the master sex determining trigger is encoded by a gene on a sex chromosome, which activates a network of downstream regulators of sex differentiation. Currently, there are more than 20 sex determining genes discovered, and as many as 13 are derived from the TGF-β pathway [20]. This pathway thus plays an important role in sex determination/differentiation of fishes but potentially regulating the number of germ cells, and/or inhibit aromatase activity to determine and maintain sex (e.g. Chen et al. [21]). Rainbow trout is a gonochoric species with an XX/XY GSD system. Interestingly, *sdY* the master sex-determinating gene of rainbow trout and other salmonid species, does not belong to the usual master sex gene families (the DM domain and Sox protein families), nor does it belong to TGF-β signaling pathway. It is a truncated duplicated copy of the immune response gene *irf9* [22]. While very specific to salmonids, the gene *sdY* is acting directly on the classical sex differentiation regulation network in all vertebrates as its protein has been shown to interact with *foxl2* [23] that is a well-known conserved female differentiation factor [24]. *foxl2* was also shown to be involved in the polled intersex syndrome in goats, where a deletion of the gene leads to triggers early testis differentiation and XX female-to-male sex reversal [25; 26].

Although phenotypic sex in rainbow trout is primarily determined by a male heterogametic GSD system, spontaneous masculinization of XX fish is a repeatedly phenomenon observed in various rainbow trout populations, at generally limited frequencies (∼1-2%) with some individuals being only partially affected (intersex) [27]. Spontaneous sex reversal is a highly heritable trait [27] and several QTL associated with this trait were detected [28, 27], highlighting the existence of several minor sex-determining genes that are independent of the major sex determinant carried by the sex chromosome. In particular, in a previous study [27], we identified in a French trout population two QTL located on chromosome 1 with a strong evidence while two suggestive QTL were identified on chromosomes 12 and 20 using the rainbow trout reference genome of Swanson line [29]. All SNP variants from these four QTL regions that covered tens of genes were kept for further investigations in the present study.

Firstly, we sought to validate the existence of the four QTL in different French rainbow trout populations, and secondly, we wanted to refine their location in the initial discovery population by combining machine learning and principal components approaches, and considering the new reference genome derived for Arlee line [30].

## Material and Methods

### Sequenced Samples

The first data set was initially produced by Fraslin et al. [27]. The 4 QTL regions that were discovered were further studied in depth in this work. The fish samples and records came from the French rainbow trout farm “Les Fils de Charles Murgat” (Beaufort, France; UE approval number FR 38 032 001). The 60 female sequenced genomes were initially aligned against the reference assembly genome Omyk_1.0 (GenBank assembly accession GCA_002163495.1) of the double haploid line Swanson from Washington State University (WSU). Following the same procedure as described in Fraslin et al. [27], we realigned these 60 sequences against the reference genome USDA_OmykA_1.1 of the WSU double haploid line Arlee (GenBank assembly accession GCA_013265735.3). This realignment was performed because French rainbow trout populations were known to be phylogenetically closer to Arlee’s genome that to Swanson’s one [31]. A total of 29,229,949 variants were obtained using a series of commands using three different variant calling tools: GATK 4.2.2.0 [32], Freebayes 1.3.5 [33] and SAMtools Mpileup 1.11 [34] Quality controls were done by using vcftools 1.15. First, indels and SNPs located on un-located contigs or mitochondrial chromosome were removed. Then, a filtering was performed on variant coverage according to the recommended settings by GATK [35]. Final numbers of 21,517,540 SNPs and 4,850,647 indels, for a total of 26,368,187 variants, were kept for further analysis. While the percentage of properly paired sequence reads over all sequenced pairs was in average 87.3% for the alignment against Swanson’s genome assembly, these statistics went up to 94.1% for the alignment against Arlee’s genome assembly.

Among the 60 sequenced females, we considered the set of 23 dams with at least 10 progeny records for sex phenotype and extreme proportions of sex-reversed offspring, either at least 25% or below 6% of neomales among their XX daughters (see S1 Supplementary Tables, S1 Table). For each of the 3 chromosomes investigated in the present study, all SNPs that exhibited polymorphism among the genomes of the 23 dams with extreme ratios of offspring sex-reversal were kept for the final analysis, i.e. 854,157 SNPs for Omy1, 965,320 SNPs for Omy12 and 492,959 SNPs for Omy20.

### Genotyped Samples

To validate the QTL detected by Fraslin et al. [27] as linked to spontaneous sex-reversal of XX trout, a set of 192 SNP were selected to design two 96 SNP genotyping arrays using microfluidic real-time PCR Fluidigm Kasp chemistry. The 192 SNPs (see S1 Supplementary Tables, S2 Table) correspond to 140 SNPs in the 2 main QTL identified on chromosome Omy1, 19 SNPs in a putative QTL region detected on Omy12 and 33 SNPs in a large 6 Mb region with putative QTL on Omy20. These arrays were used to genotype 315 fish from 6 diverse French populations of rainbow trout named by the letters A to F (Table 1) with at least 30 XX fish, including a minimum of 8 neomales, per population. Population A was made up of XX sibs from the same birth cohort as the parents used to produce the QTL discovery population [27]. Respectively, fish from populations B and C and those from populations E and F came from two other French breeding compagnies, while fish from population D came from a commercial site using fry from unknown origin.

**Table 1.** Description of the 6 rainbow trout XX populations used for QTL validation

| Population | A | B | C | D | E | F |
| --- | --- | --- | --- | --- | --- | --- |
| Number of genotyped fish | 49 | 30 | 64 | 77 | 45 | 50 |
| Phenotyped as female | 34 | 22 | 32 | 29 | 31 | 32 |
| Phenotyped as male (including intersex) | 15 | 8 | 32 | 48 | 14 | 18 |
| Phenotyped as pure male (no intersex) | 8 | nr | 17 | 32 | 6 | nr |

nr: values not reported by the breeding compagny

Pieces of caudal fin sampled from those 315 fish were sent to Gentyane genotyping platform (INRAE, Clermont-Ferrand, France) for genotyping after DNA extraction using the DNA advance kit from Beckman Coulter following manufacturer instructions.

After quality control, 19 SNPs were eliminated (including 14 from Omy1) from the analyzes of all the populations because their genotyping rate was below 90%.

### Statistical analysis for validation of the existence of the QTL

An exact Fisher test was run for each of the 6 validation populations using the fisher.test function in R version 4.3.1 [36] which implements the method developed by Mehta and Patel [37, 38] and improved by Clarkson, Fan and Joe [39] to test the independence of genotypes at each SNP and phenotypic sex of fish.

This test was applied for the set of 173 validated SNPs across all six populations. In any of these populations, ∼170 SNPs were polymorph and had a genotyping rate above 90%. To account for multiple testing using Bonferroni correction, we considered that SNPs were significant at the genome level in a given population when p_value < 0.0005 (i.e. log_10_(p_value) < 3.3, assuming that about 100 SNPs were not highly redundant among ∼170 tested for each population) and had only a putative effect when p_value < 0.005 (i.e. log_10_(p_value) < 2.3).

To confront the results between the validation and discovery populations, the latter threshold value was also used to detect, by exact Fisher test, the association of all the sequence variants observed in the 4 QTL regions with the average progeny sex-ratio of the 23 extreme dams of the discovery population. A second, more stringent, threshold (p_value < 0.002) was considered to focus on the main significant associations with sequence variants in the discovery population. Because of the strong linkage disequilibrium observed across SNPs along the chromosome [40], the QTL region on Omy20 was strongly expanded to test, one-by-one, all the variants spanning the chromosome from 27 to 38 Mb.

### Statistical approaches for refinement of QTL locations in the discovery population

#### Machine learning approach using Random Forests

Random forest (RF) is a machine learning method that aggregates complementary information from an ensemble of classification or regression trees trained on different bootstrap samples (animals) drawn with replacement from the original data set [41]. First, every tree is built using a bootstrap sample of the observations. Second, at each node, a random subset of all predictors (the size of which is referred to as mtry hereafter) is chosen to determine the best split rather than the full set. Therefore, all trees in a forest are different. A particularity of the RF is the out-of-bag data, which corresponds to the animals not included (roughly 1/3) in the bootstrap sampling for building a specific tree. It can be used as an internal validation set for each tree, which allows the computation of RF error rate based on misclassified animals in the out-of-bag data. RF can be applied successfully to “large p, small n” problems, and it is also able to capture correlation as well as interactions among independent variables [42].

We considered the five following genomic zones to define haplotypes of subsequent SNPs spanning from 67 to 69Mb on Omy1, from 8.5 to 9.5Mb on Omy12, and, on Omy20: 27-29Mb, 33.5-35.5 Mb, and 36-37Mb. In total, we considered 21,310 variants for Omy1, 12,191 for Omy12, and 56,643 for Omy20, whatever their distance to genes in the QTL regions.

We recoded the genotypes depending of the number of reference alleles present in the genotype: “2” for a homozygous reference genotype, “1” for a heterozygous genotype, “0” for the alternative homozygote; missing genotypes were encoded “5”.

To minimize the RF error rates, we ran a preliminary set of analyses to establish the optimum SNP window size (swind), the number of sampled trees (ntree), and the number of sampled predictor variables (mtry) chosen at random to split each node. For any analysis procedure, the more highly correlated the variables are, the more they can serve as surrogates for each other, weakening the evidence for association for any single correlated variable to the outcome if all are included in the same model. Therefore, to avoid redundancy of information and limit the number of variables to analyze, we built some haplotypic genotypes by retaining one SNP out of two and gathering them in the same haplotype across a window of 80 initial SNPs (correspond to swind=40). We varied swind from 20 to 200 subsequent SNPs, ntree from 100 to 1,000 and mtry from 5 to 20. The final parameter set (swind=40, ntree=400 and mtry=7) was chosen as consistently giving the lower median error rate over 50 runs for any of the three tested chromosomes. A final set of 246 haplotypes for Omy1 (131 for Omy1_a and 115 for Omy1_b), 152 haplotypes for Omy12 and 707 haplotypes for Omy20 (247, 288, and 172, respectively for Omy20_a, Omy20_b and Omy20_c) were then analyzed by averaging results across 100 RF runs. We used the package RandomForest 4.18 [43] in R version 4.3.1 [36].

Haplotypes were ranked by decreasing order of importance in the prediction model, using the mean decrease in Gini index. The Gini index measures the probability for a specific feature of being misclassified when chosen randomly. The higher is the value of mean decrease in Gini index, the higher is the importance of the variable in the model.

A list of 45 positional candidate genes was established based on results of the best ranked haplotypes as well as the preliminary Fisher exact tests on SNP.

#### Discriminant Analysis of Principal Components to identify the most promising variants

The Discriminant Analysis of Principal Components (DAPC) helps explore the genetic structure of biological populations. This multivariate method consists in a two-step procedure: 1) genetic data are centred and used in a Principal Component Analysis (PCA); 2) a Linear Discriminant Analysis is applied to the principal components of PCA. A basic matrix operation is used to express discriminant functions as linear combination of alleles, allowing one to compute allele contributions, i.e. to identify the most interesting variants [44].

We considered the 33,731 SNPs that were in the list of genes identified by RF or Fisher’s exact test. We also studied the SNPs in the close upstream and downstream regions of theses genes (±2 kb). Filtering out SNPs with call rate below 100% using PLINK 1.9 [45], we applied DAPC on a final set of 27,828 variants without any missing genotypes. We used the R adegenet package [46] to find by DAPC the main variants that best discriminate between dams with high and dams with low sex-reversal ratios in their offspring.

### Annotation of genes and variants

Genes within QTL regions were annotated using the NCBI *O. mykiss* Arlee genome assembly USDA_OmykA_1.1. (GCA_013265735.3) (Gao et al. 2021). In addition to NCBI gene summaries, functional information for genes was extracted from the human gene database GeneCards® (https://www.genecards.org/) that also includes protein summaries from UniProtKB/Swiss-Prot (https://www.uniprot.org/uniprotkb/).

SNP annotation was performed for the 33,731 SNPs identified for the list of genes of interest, applying the software SNPEff4.3T [47] on the NCBI annotation file annotation released for *O. mykiss* Arlee genome reference assembly (GCA_013265735.3). These annotations permitted to retrieve any information about the predicted impact of the variants on the genes, and, in some cases, on the resulting proteins. Interesting variants were further studied using the Genome Data Viewer of NCBI to indicate their intronic or exonic positions within the genes.

## Results

### Validation of the existence of the 4 QTL in various French rainbow trout populations

Among the 173 SNPs that could be tested for their association with spontaneous sex-reversal in the validation populations, 117 had a significant effect in at least one population. Among the latter, 50 SNPs on Omy1 had at least an effect in population A that was genetically close to the QTL discovery population. Most of these 50 SNPs had also an effect observed in populations B, C and E. On the contrary, none of the SNPs tested on Omy12 or Omy20 had an effect in population A, probably because of sampling issues for low frequency variants in the population. However, 7 SNPs on Omy12 (out of 17 that could be tested) had an effect in populations C, D, E and F; and 13 SNPs on Omy20 (out of 30 that could be tested) had an effect in populations C, D and F.

The list of the 90 SNPs that were at least associated with a putative effect in 2 validations populations are given in S1 Supplementary Tables, S3 Table with indication of the level of significance in those populations. Table 2 reported the 63 SNPs that had at least a putative effect in four validation populations, or a clear significant effect in at least two populations. Out of these 63 SNPs, 50 were located within genes or in their close vicinity (< 5kb) and NCBI annotation for those genes are then reported in Table 2.

**Table 2.**
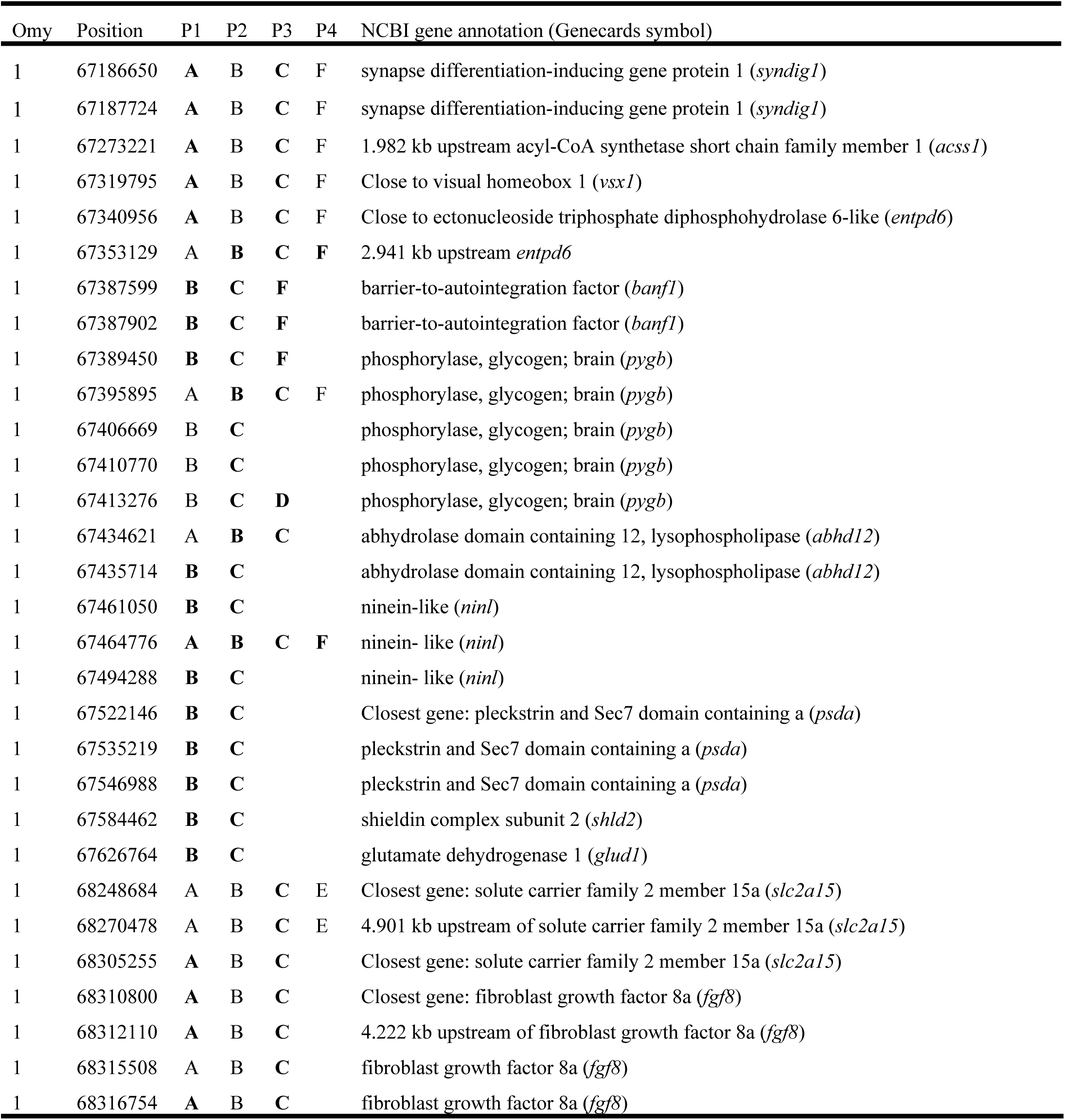
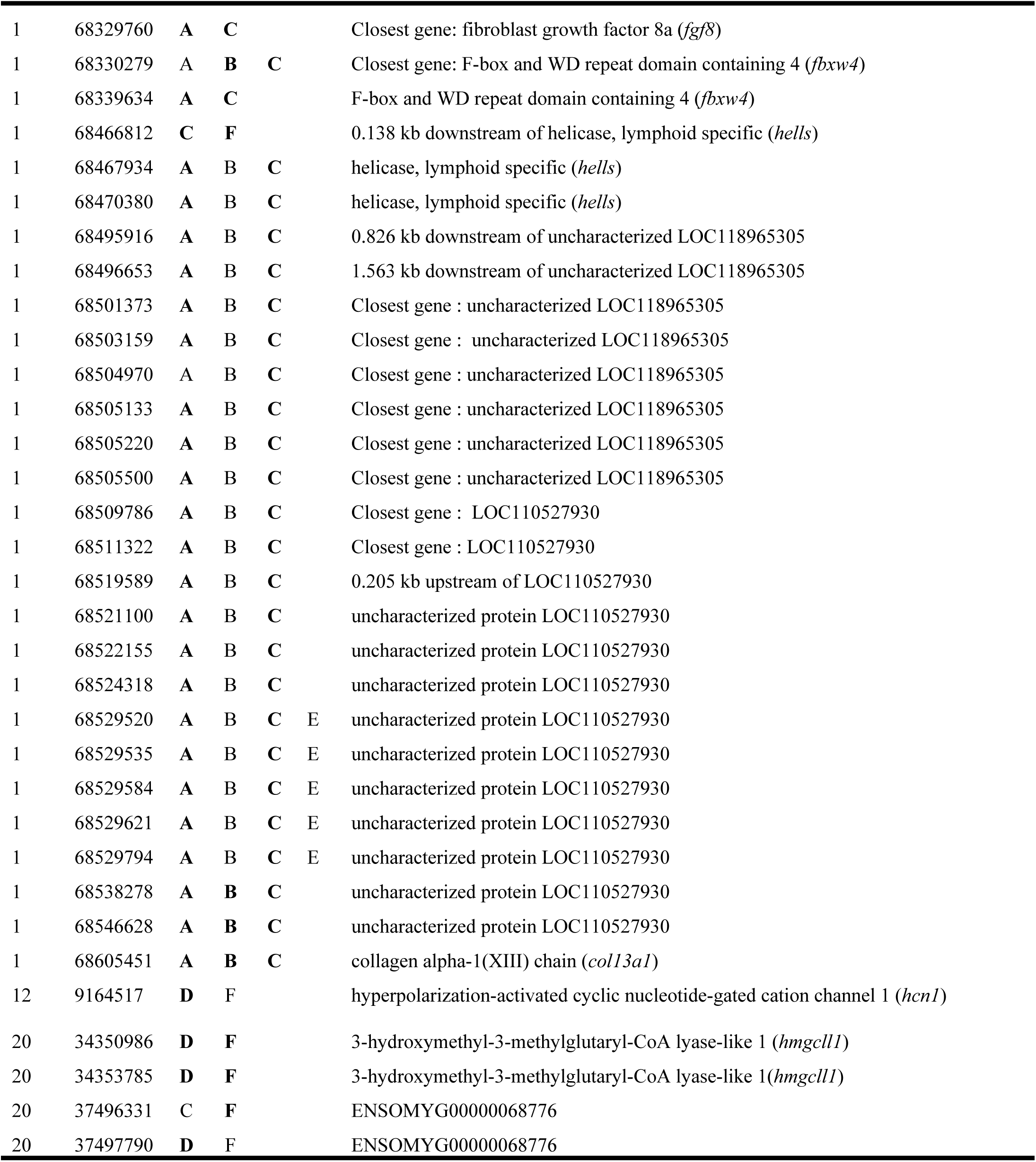
Most significant SNPs in at least 2 out of the 6 validation populations named A to F (names in bold correspond to a p_value < 0.0005, others to a p_value < 0.005)

### Extension of the QTL region boundaries in the discovery population

Because most of the SNPs located at the boundaries of the QTL regions reported by Fraslin et al. [27] had a significant effect in some of the validation populations, we applied Fisher’s exact test on the dams’ average progeny phenotypes in the discovery population for all sequence variants located in QTL regions enlarged by 0.5 Mb on each side. Based on these tests, we defined the new extended boundaries for the QTL that are given in Table 3. In particular, we enlarged the search of QTL in 3 different regions of Omy20, spanning from 27 to 29 Mb, 33.5 to 35.5 Mb, and 36 to 38 Mb, respectively for Omy20_a, Omy20_b and Omy20_c.

**Table 3.**
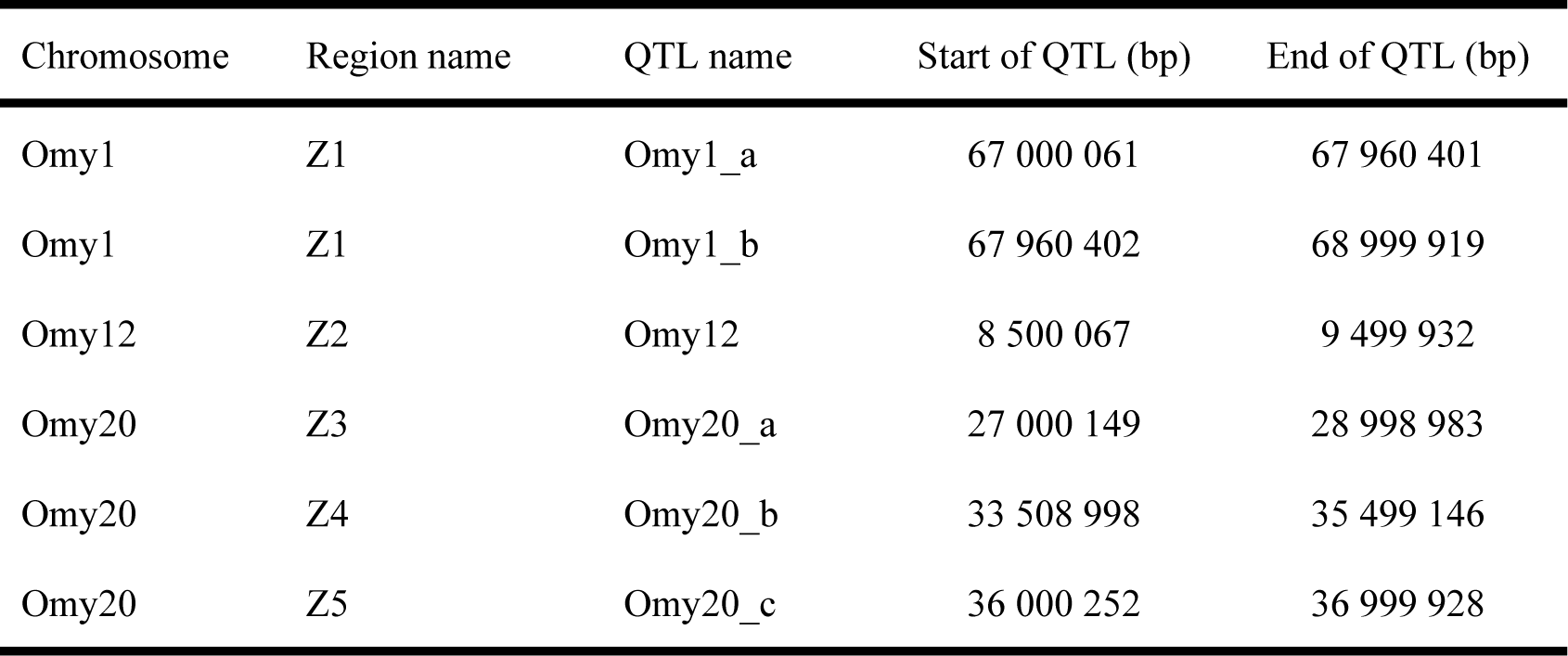
Extended boundaries of sex-reversal QTL regions in the current study (location on Arlee reference genome) compared to QTL intervals reported by Fraslin et al. [27]

| Chromosome | Region name | QTL name | Start of QTL (bp) | End of QTL (bp) |
| --- | --- | --- | --- | --- |
| Omy1 | Z1 | Omy1_a | 67 000 061 | 67 960 401 |
| Omy1 | Z1 | Omy1_b | 67 960 402 | 68 999 919 |
| Omy12 | Z2 | Omy12 | 8 500 067 | 9 499 932 |
| Omy20 | Z3 | Omy20_a | 27 000 149 | 28 998 983 |
| Omy20 | Z4 | Omy20_b | 33 508 998 | 35 499 146 |
| Omy20 | Z5 | Omy20_c | 36 000 252 | 36 999 928 |

In total, 246 SNPs had a p_value ≤ 0.005 (see S1 Supplementary Tables, S4 Table), and the 19 SNPs with a more stringent p_value ≤ 0.002 were mainly located within genes or in their close vicinity (Table 4).

**Table 4.**
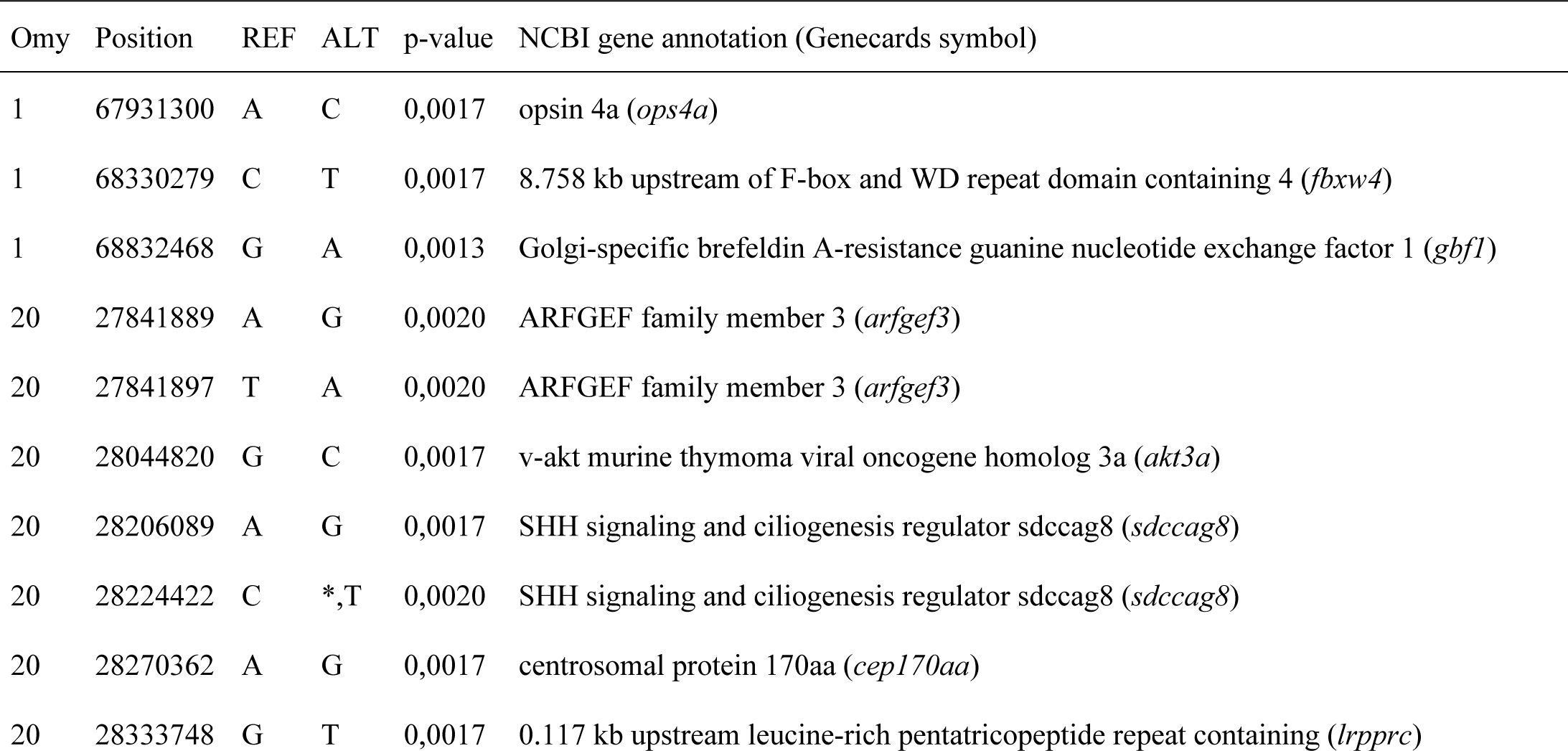
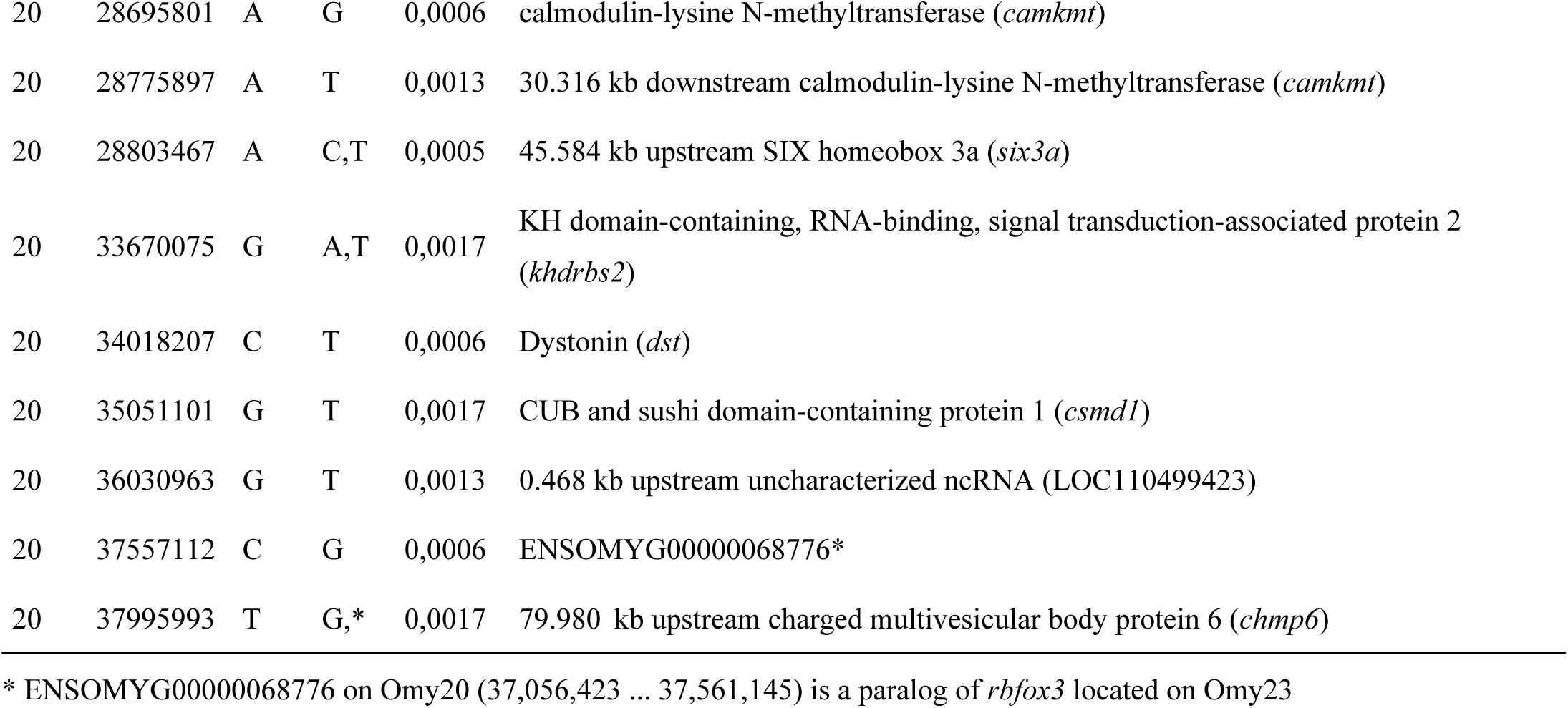
Most significant SNPs by Fisher’s exact test (p_value ≤ 0.002) considering the 23 dams with extreme progeny sex-ratios in the QTL discovery population.

### Identification of the most relevant haplotypes and genes explaining sex-reversal in the discovery population

Initially, we ran a RF analysis for each of the 3 chromosomes with all their QTL regions jointly analyzed. Respectively for the RF on Omy1, Omy12 and Omy20, we presented the 30, 15 and 30 best ranked haplotypes based on Gini index as well as the corresponding genes, in S5, S6 and S7 Tables. For the 2-Mb region on Omy1 (246 haplotypes spanning from 67 to 69 Mb), the median error rate was 28% with a minimum error rate of 15% and a maximum rate of 37% of fish phenotypes wrongly classified as the opposite sex across 100 RF simulations. While the RF analysis of error rate indicated that the 2 Mb tested on Omy1 were really useful to discriminate the dams with high and low progeny sex-reversal ratios, the median error rate derived from the RF analysis on Omy12 (152 haplotypes spanning 8.5 to 9.5 Mb) was 42% (ranging from 29% to 51% across the 100 RF runs), indicating that the best haplotypes and genes identified on this chromosome were not so strongly associated with sex-reversal. In terms of error rate, the RF analysis on the 707 haplotypes covering the QTL regions on Omy20 (247 haplotypes from 27 to 29 Mb, 288 haplotypes from 33.5 to 35.5 Mb, and 172 haplotypes from 36 to 37 Mb) gave intermediate results in-between the two previous analyses, misclassifying one third of the dams on average with a minimum error rate of 29% and a maximum rate of 38% across 100 RF simulations.

These results are consistent with the higher proportions of genetic variance explained by QTL we estimated on Omy1 than on Omy12 or Omy20 [27]. In addition, we considered all together the 1,105 haplotypes covering the QTL regions along the three chromosomes in a last RF analysis. The empirical distribution of error rates for the full RF analysis gave median, minimum and maximum values similar to the ones of the RF on Omy20 (33%, 29%, and 38%, respectively) while we were expecting that these values will be at least as low as for the RF on Omy1. Due to the very small size of our dataset, it is likely that we had too many variables in the full RF analysis, creating some « noise » by considering over 1,100 haplotypes together in the analysis to classify our 23 individuals.

Looking at the locations (Fig.1) of the best ten RF ranked haplotypes on the 2 Mb evaluated on Omy1 (see S1 Supplementary Tables, S5 Table), they fell within the nine following genes, six of them being identified with significant effects on sex-reversal in several validation populations (Table 2) or within some significant SNPs in the discovery population (Table 4): *sdhaf4*, *tlx1*, *syndig1* and *vsx1* genes located in QTL Omy1_a; *cep68*, *fbxw4*, *hells*, LOC110527930, and *gbf1* in QTL Omy1_b. The first best-ranked haplotype included two genes from QTL Omy1_b (*sdhaf4* and *tlx1*), while the second and third best-ranked haplotypes were in the sequence of the *syndig1* gene on Omy1_a.

**Fig. 1.**
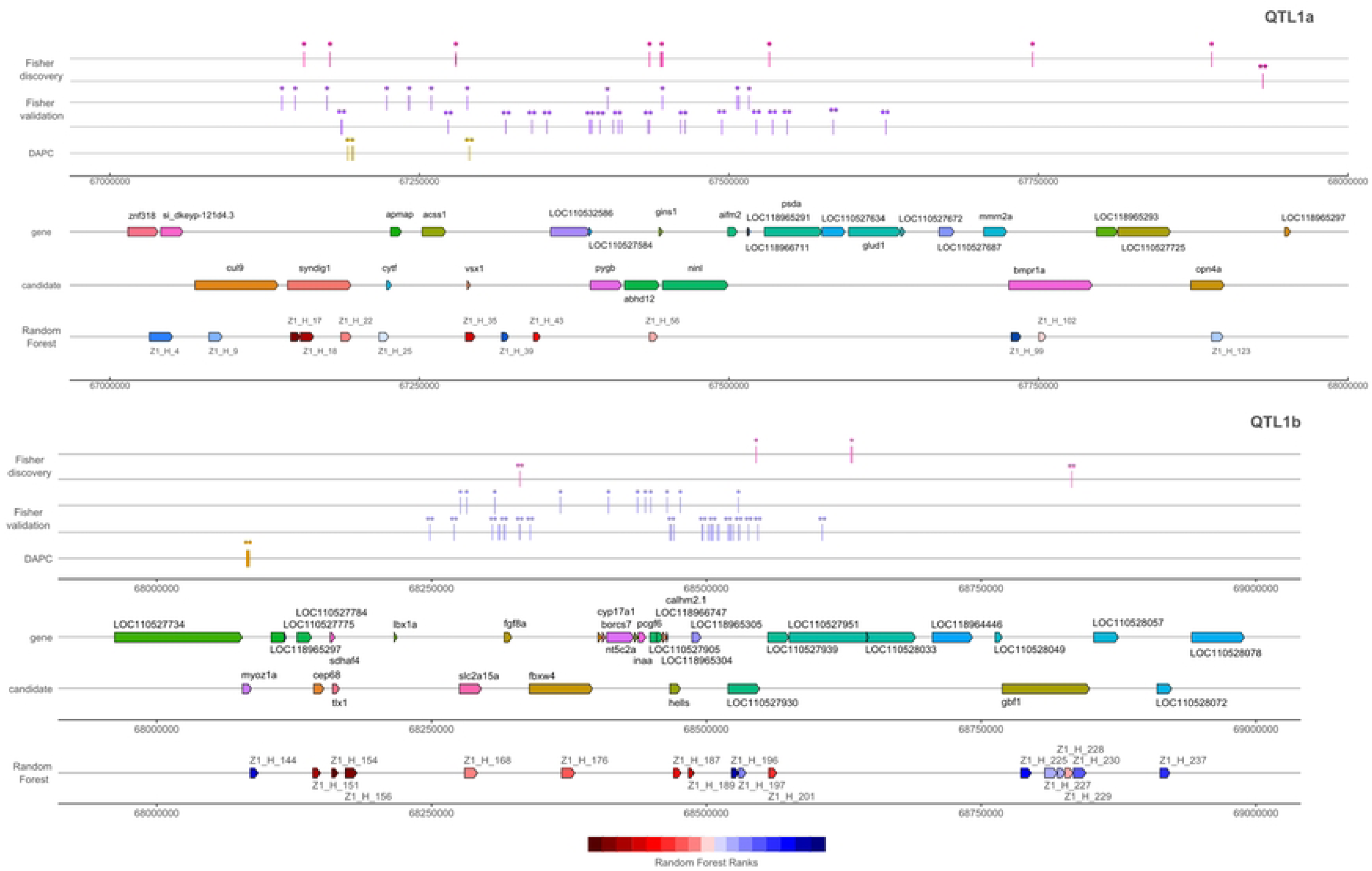
Positions of genes within QTL regions a and b on Omy1 with significant SNPs positions in discovery or validation population as well as best ranked haplotypes

These results are not fully consistent with our first GWAS [27] using progeny imputed genotypes and Bayesian mixed model methodologies. In this previous analysis, the *syndig1* gene was not included in the credibility interval for Omy1_a. Although there was no SNP within this gene among the most significant SNPs (p_value ≤ 0.002) in the discovery population, 2 SNPs had, however, a p_value < 0.005 (see S1 Supplementary Tables, S4 Table). In addition, *syndig1* exhibited highly significant SNPs in four validation populations, including population A. All these results make *syndig1* a very plausible candidate gene for QTL Omy1_a. As regards to the initial QTL Omy1_b, there is now an extended credibility interval, including genes before *slc2a15a* such as *tlx1* and *cep68*, and genes after the uncharactezized LOC110527930 such as *gbf1*. While LOC110527930 initially was our favourite causative gene for Omy1_b, genes such as *tlx1*, *cep68*, *fbxw4* and *hells* are now prefered within Omy1_b region (Fig. 3).

On the 1 Mb tested on Omy12 (see S1 Supplementary Tables, S6 Table), the first three best haplotypes to explain sex-reversal were within the same two *hcn1* and *ccbe1* genes that were also identified in three populations unrelated to the discovery population (see S1 Supplementary Tables, S3 Table).

On the 5 Mb tested on Omy20 (Fig.2), unfortunately, there was no variant tested in the validation populations for the eight genes corresponding to the ten best ranked haplotypes under RF analysis (see S1 Supplementary Tables, S7 Table) that were in decreasing order of importance: *lrrc59*, *csmd1*, *khdrbs2*, *abcg5*, *akt3*, *rgs17*, *caskin2* and *dst*. However, some highly significants SNPs are detected in the discovery population for four of these genes (*akt3*, *khdrbs2*, *dst* and *csmd1*, see Table 4).

**Fig. 2.**
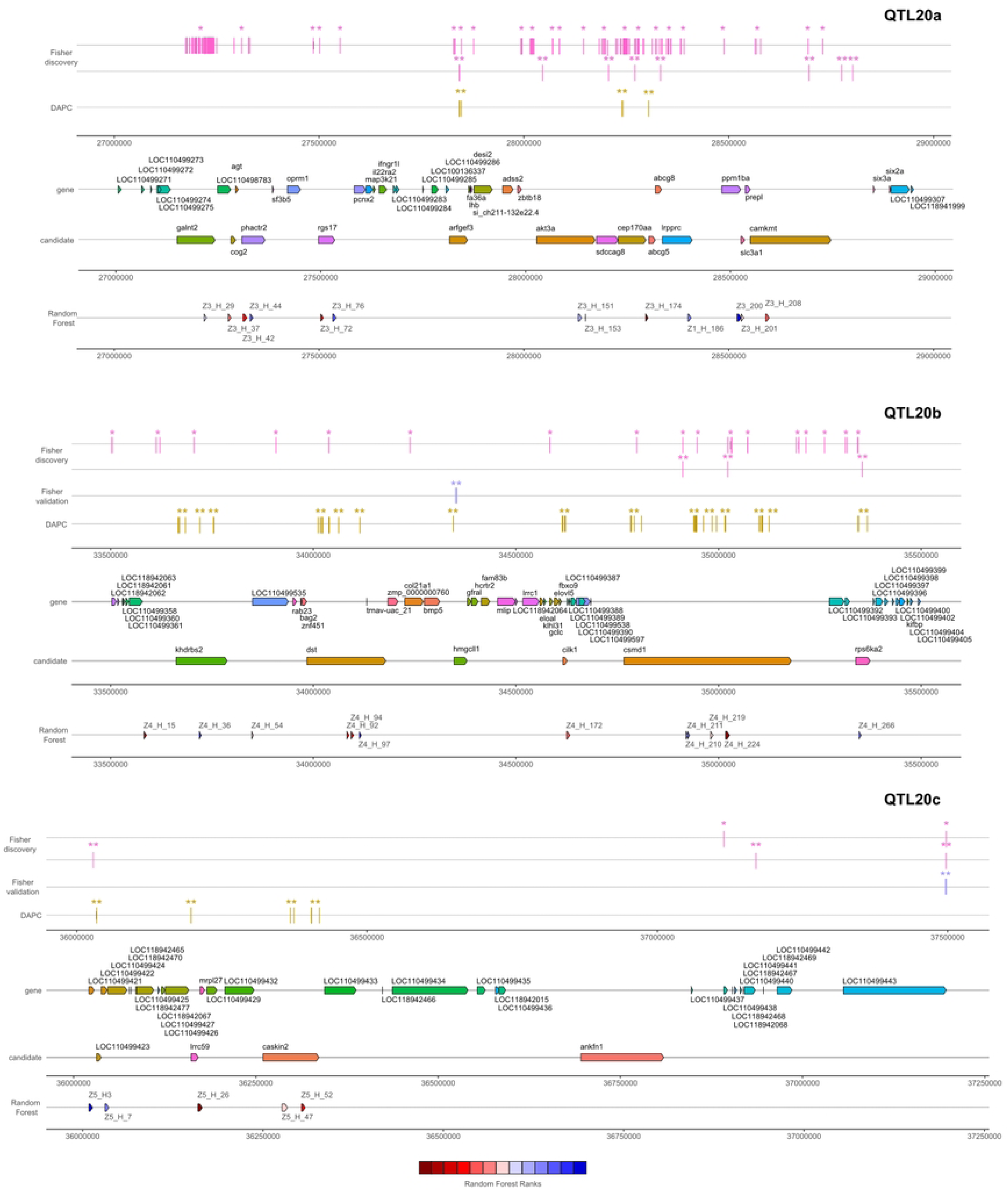
Positions of genes within QTL regions a, b and c on Omy20 with significant SNPs positions in discovery or validation population as well as best ranked haplotypes

As regards to the ranking of the best haplotypes to explain sex-reversal across all the QTL regions spanning the 3 chromosomes (see S1 Supplementary Tables, S8 Table), the first one was positionned within *lrrc59* gene on Omy20_c, the second and third ones included *sdhaf4* and *tlx1* on Omy1_b, and the fourth one was located within the *csmd1* gene on Omy20_b. The next four best-ranked haplotypes were all on Omy20, located within *abcg5*, *caskin2*, *khdrbs2*, and *dst* genes. Some genes identified on Omy1 in validation populations (*syndig1*, *hells*, *fbxw4* and LOC110527930) appeared then as the subsequent haplotypes ranked with very similar Gini indices (Fig. 1). Noticeably, most of the best-ranked haplotypes concerned sequences within genes, and were consistently ranked between the full RF analysis and the RF limited to their chromosomic location.

Therefore, based on the results of the 60 best ranked haplotypes on the Gini index of this full RF analysis, as well as the additional results from Fisher’s exact tests (Tables 2 and 4), we identified a list of 45 genes (see S1 Supplementary Tables, S9 Table) whose variants in their sequence or in their close vicinity (2kb) were the most likely to explain a significant part of sex-reversal phenotypes.

### Identification of the potential causative variants combining DAPC and SNP annotations

We considered the corresponding sequence variants of those genes (±2 kb) in a DAPC in order to find the best 100 SNPs to discriminate among the 23 dams the ones with high progeny sex-reversal ratios from those with low ratios. Running a simple ACP on all those variants do not allow to correctly discriminate the two groups of dams as shown by Fig.3A. On the contrary, the DAPC allowed us to perfectly discriminate the two groups as demonstrated in Fig.3B.

**Fig. 3.**
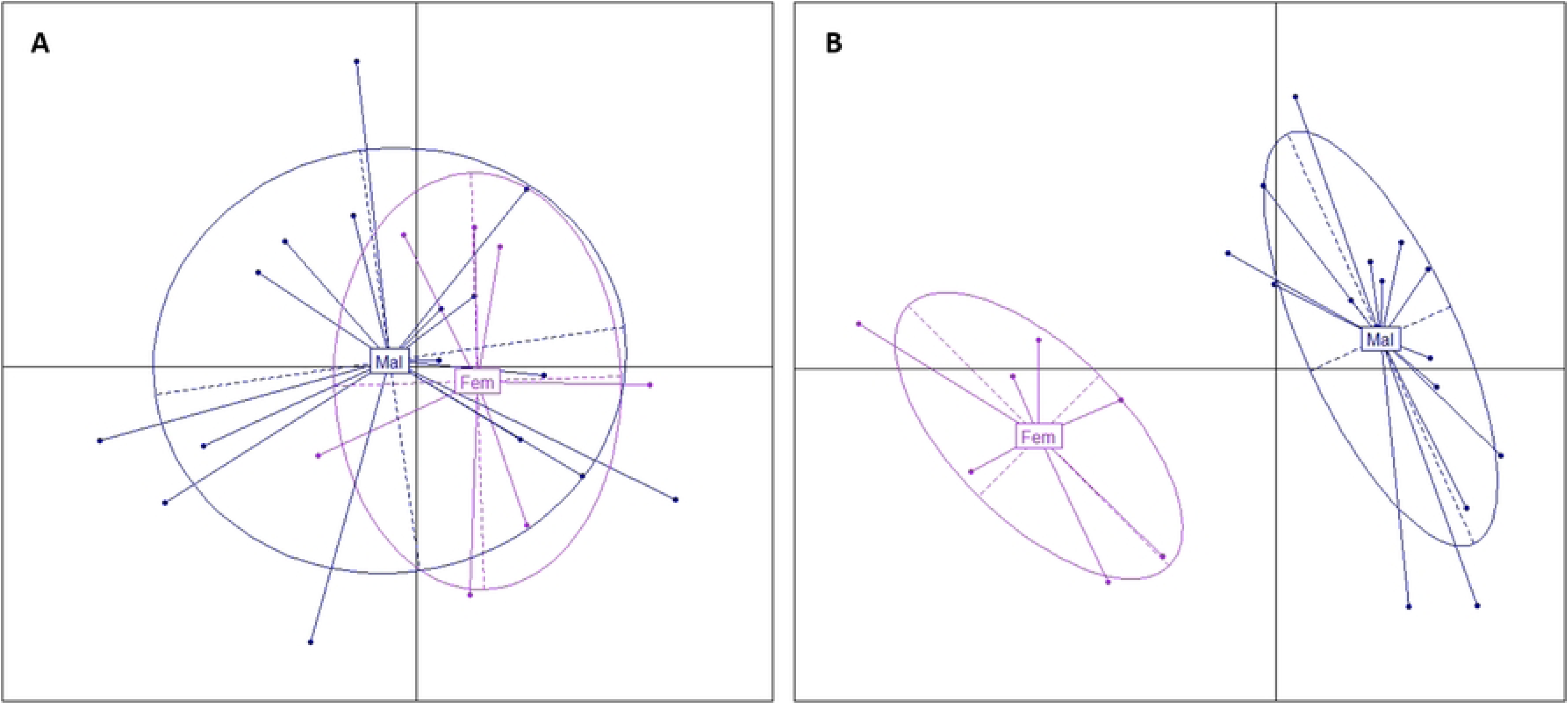
PCA of variants between “Fem” and “Mal” dam groups on 45 candidate genes. **A. PCA of the variants in the 45 candidate genes (±2 kb) and B. PCA of the 100 best variants selected by DAPC to discriminate the “Fem” group of the seven dams with low progeny sex-reversal ratios and the “Mal” group of the 16 dams with high progeny sex-reversal ratios.**

In S1 Supplementary Tables, S10 Table are given the genotypes for the 100 best SNPs that were genotyped for the 23 dams among 27, 828 SNPs (without missing genotypes out of 33,731 SNPs), as well as for 5 additional dams with phenotypic sex recorded on at least 6 offspring in the discovery population (see S1 Supplementary Tables, S1 Table). We considered these 5 dams as an internal validation: 3 dams (noted AS, AT and AU) had a high progeny sex-reversal ratio while no sex-reversed offspring were observed for the last 2 dams (noted BS and BT). Those best 100 SNPs were located within or nearby the downstream or upstream regions of 16 genes belonging to the list of the 45 genes previously identified by RF and Fisher’s exact test: on Omy1, *syndig1*, *vsx1*, and *myoz1*; on Omy12, *hcn1* and *ccbe1*; on Omy20, *arfgef3*, *cep170aa*, *abcg5*, *khdrbs2*, *dst*, *hmgcll1*, *cilk1*, *rps6ka2*, LOC110499423, *lrrc59* and *caskin2*. Most of these 100 SNPs were located in intronic regions of the genes, two were synonymous mutations and none were splice donor variants in introns or missense variants in exons. When focusing on the 50 best SNPs to discriminate between the two groups of dams, only *hcn1* remained on Omy12; *syndig1* and *myoz1* were kept for Omy1_a and Omy1_b, respectively; and, all genes, except LOC110499423, remained for QTL regions on Omy20.

## Discussion

### Significance of the different QTL regions across French rainbow trout populations

Considering the list of the 100 best-ranked SNPs under DAPC in the discovery population confirms the hypothesis of two QTL on Omy1, one QTL on Omy12, and lead us to suggest the existence of three distinct QTL on Omy20. All the QTL regions previously identified [27] are confirmed as playing a role on sex-reversal in at least two populations different from the discovery one. The most significant SNPs across several populations are located within genes or in their close vicinity.

Regarding QTL Omy1_a, the candidate genes that appear as the most relevant to explain sex-reversal across populations were *syndig1*, *acss1*, *vsx1*, *entpd6*, *pygb* and *ninl*. In the initial study [27], the peak SNPs for this QTL were located either within the *pygb* or *ninl* gene, depending on the statistical approach. In particular, Fraslin et al. [27] mentioned a SNP located at 63,543,061 bp (on Swanson reference genome) within the *ninl* gene and annotated has a missense variant. This SNP was not genotyped in the validation population, but in this study, a very close SNP located at 63,530,416 bp on Swanson reference genome (corresponding to 67,434,621 bp on Arlee reference genome) has a putative effect in population A and a significant effect at the genome level in populations B and C (see S1 Supplementary Tables, S3 Table).

As regards to the main QTL Omy1_b, Fraslin et al. [27] identified it with all peak SNPs located in the intergenic region between the *hells* gene and the uncharacterized protein LOC110527930, the latter being proposed as the most relevant positional candidate for explaining 14% of the genetic variance of sex-reversal in the discovery population. However, in the present study, other genes should also be considered as candidate genes in several validation populations. Some were located before the *hells* gene (*slc2a15a*, *fgf8*, and *fbxw4*), while the most distal one, *col13a1*, was located after LOC110527930. This observation, as well as the identification of *syndig1* in QTL Omy1_a that was positioned before the first QTL interval defined in Fraslin et al. [27] were the motivation to further study the sequence variants in larger genomic regions.

In the previous study [27], the QTL on Omy12 was detected using the 57K chip, but not considering imputed sequence variants in the discovery population. In the present study, this QTL is not validated in population A although this population is closely related to the discovery one. Nevertheless, the QTL is validated in population D and F with a significant SNP identified within *hcn1* gene. We can hypothezise that the QTL is then segregating in the discovery population, but either at a too low frequency or too small effect to be statistically detected.

The QTL region on Omy20 is not validated in population A, but two QTL are segregated on Omy20 in populations C, D and F, with significant SNPs located in two distant genes: *hmgcll1* located around 34.4 Mb, and ENSOMYG00000068776 gene located around 37.5 Mb. The same hypothesis as for the QTL on Omy12 can thus be proposed for the absence of statistical validation in population A. In addition, in an INRAE experimental rainbow trout line, a QTL for spontaneous sex-reversal was also identified on Omy20 (previously named RT17) but with a very imprecise position [28].

In the next paragraphs, we describe the putative functional roles of our best positional candidate genes for the two QTL on Omy1 as well as for the most promising genes for the three different QTL on Omy20. Other positional candidate genes on Omy12, Omy20_a and Omy20_b with less functional evidence are presented in an additional section [see S1 Appendix].

### *syndig1* as the best positional and functional candidate gene for QTL Omy1_a

It is very likely that one or a few variants in the *syndig1* (synapse differentiation-inducing gene protein 1) sequence or in its close downstream region (< 1kb) are the causative variants explaining a part of the sex-reversal phenotype. First of all, 4 SNPs were significant in two to four validation populations (see S1 Supplementary Tables, S3 Table). In the discovery population, the RF analysis highlighted 3 haplotypes in the gene (see S1 Supplementary Tables, Suplementary table 5), among which 3 missense variants were annotated. The DAPC ranked 3 SNPs within or dowstream *syndig1* (see S1 Supplementary Tables, S10 Table) among the 100 best SNPs to discriminate the low and high sex-reversed progeny dams. However, among the 50 best SNPs, the only 2 retained were located in downstream intergenic positions. We speculate that these latter positions were kept to the detriment of SNPs within *syndig1* due to linkage desequilibrium between SNPs of the two close genes *syndig1* and *vsx1* (visual system homeobox 1). Indeed, a unique SNP was retained from *vsx1* in the list of the 100 best SNPs and was filtered out in the list of the 50 best ones. We may speculate that *vsx1* may play a minor additional role to explain sex-reversal as significant SNPs were detected both within gene in the discovery population and in an upstream region in 4 validation populations. It is worth mentionning the existence of 5 missense variants in *vsx1*. This gene is expressed during early embryogenesis as well as in the adult retina. It participates in regulating retinal progenitor proliferation, differentiation and functional maintenance of bipolar cells [48, 49, 50]. As far as we know, no direct link has been suggested between *vsx1* and sex determination in any species.

**Table 5.** Protein sequence identities of genes associated to sex-reversal in rainbow trout across six species in Vertebrates

| rainbow trout /<br>target<br>(% id AA) | Atlantic<br>salmon | Medaka | Zebrafish | Mouse | Sheep | Human |
| --- | --- | --- | --- | --- | --- | --- |
| *syndig1* | 98% | 58% | 69% | 65% | 67% | 59% |
| *vsx1* | 96% | 69% | 84% | 62% | 47% | 52% |
| *fbxw4* | 99% | 68% | 71% | 58% | 58% | 58% |
| *hells* | 98% | 79% | 79% | 65% | 64% | 64% |
| *myoz1a* | 92% | 56% | 63% | 39% | 37% | 40% |
| *tlx1* | 95% | 75% | 82% | 61% | 58% | 61% |
| *gbf1* | 99% | 82% | 83% | 72% | 72% | 72% |
| *hcn1* | 99% | 84% | 84% | 74% | 68% | 68% |
| *arfgef3* | 99% | 76% | 84% | 71% | 72% | 72% |
| *cep170aa* | 98% | 63% | 68% | 53% | 54% | 55% |
| *abcg5* | 100% | 91% | 94% | 87% | 90% | 88% |
| *khdrbs2* | 98% | 76% | 82% | 70% | 69% | 69% |
| *dst* | 96% | 72% | 71% | 60% | 59% | 62% |
| *cilk1* | 99% | 70% | NA | 32% | 55% | 55% |
| *csmd1* | 99% | 79% | 89% | 76% | 76% | 77% |
| *lrrc59* | 98% | 71% | 62% | 58% | 58% | 58% |
| *caskin2* | 91% | 54% | 54% | 59% | 59% | 59% |

As regards to *syndig1*, this gene encodes a protein that belongs to the interferon-induced transmembrane family of proteins. It plays a critical role during synapse development to regulate *ampar* (α-amino-3-hydroxy-5-methyl-4-isoxazolepropionic acid receptor) content and have a role in postsynaptic development and maturation [51, 52]. Although *syndig1* expression is mainly restricted in the embryonic retina in mice [53], its expression has been detected also in granulosa cells at the small antral stage in sheep [54] suggesting a potential role in follicular development in vertebrates.

### Multiple candidate genes for QTL Omy1_b, with two favourites: *tlx1* and *hells*

While this QTL was identified as the one explaining the largest proportion of genetic variance of spontaneous sex-reversal [27], the previous study pointed on different candidate genes than in the current one. In the first study, the three putative functional candidate genes were *fgf8a*, *cyp17a1* and LOC110527930 based on progeny imputed genotypes, while the current analysis of dams’ sequences pointed to genes located upstream, prioritizing *tlx1* (T cell leukemia homeobox 1) for RF analysis and *myoz1* (myozenin-1) for DAPC. It seems, in fact, that two different QTL should be distinguished within Omy1_b, Omy1_b1 and Omy1_b2, on each side of *cyp17a1*. Although *cyp17a1* is known to be involved in gonad masculinization in the common carp and zebrafish [55, 56], no variant in this gene was significantly associated with sex-reversal in rainbow trout, neither in the discovery population, nor in the validation ones. The new analyses highlighted genes, *myoz1* and *tlx1*, that are located in region Omy1_b1 before the start of the QTL region previously defined at *slc2a15a* gene [27]. However, it is possible that a cis-regulatory element within the QTL region could control at long-distance the expression of *cyp17a1*, influencing sex differentiation in rainbow trout. Such long-distance cis-regulatory elements have been found to control sex like in the Sry-deficient Amami spiny rat where the male-specific upregulation of *sox9* is controlled by an enhancer region 0.5 Mb upstream of *sox9* [57].

The initial Omy1_b is now defined as a second genomic region Omy1_b2, spanning from *fbxw4* to *gbf1* (including *hells* and LOC110527930). While the region Omy1_b2 was clearly associated with sex-reversal in four validation populations, there was no SNP among the 100 best ranked by DAPC. The RF analysis, however, identified some haplotypes among the 10 best ones to explain sex-reversal on Omy1, putting similar emphasis to four genes: *fbxw4*, *hells*, LOC110527930 and *gbf1*. In addition, Fisher’s exact test gave a p_value ≤0.002 for a SNP within *gbf1* (Golgi-specific brefeldin A-resistance guanine nucleotide exchange factor 1) in the discovery population, while SNPs in this gene were unfortunately not tested in the validation populations as outside the limits of the initial credibility interval for the QTL. As far as *fbxw4* (F-box and WD repeat domain containing 4) gene is concerned, no direct link with gonad development, oogonia or spermatogognia could be established. This gene is a member of the F-box/WD-40 gene family that may participate in Wnt signaling which is a pathway involved in gonadal differentiation (see S1 Supplementary Tables, S9 Table). As regards to *gbf1*, it is one of the guanine-nucleotide exchange factors (GEF) for members of the Arf family of small GTPases involved in vesicular trafficking in the early secretory pathway (see S1 Supplementary Tables, S9 Table). A *gbf1* expression was shown throughout spermatocyte development in the Golgi apparatus with expression extending into early spermatids where it localizes also to acrosome [58]. The positionnal candidate gene *hells* (helicase lymphoid specific) in Omy1_b2 is a very convincing functional candidate gene for sex-reversal. *hells* gene (also known as *lsh*) is a member of the *snf2* helicase family that is involved in cellular proliferation, and required for normal development and survival in the mouse [59]. s*nf2*-like helicases play a role in chromatin remodeling, and the *hells* protein has been implicated in the control of genome-wide DNA methylation [60]. Analysis of ovarian explants obtained from *hells* mutant females demonstrates that lack of *hells* function is associated with severe oocyte loss and lack of ovarian follicle formation [61]. Although *hells* is expressed in undifferentiated embryonic stem cells, it is significantly downregulated upon differentiation [62]. Because of the neonatal lethality of the *hells* -/- mice, Zeng et al. [63] used allografting of testis tissue from *hells* -/- mice to study postnatal male germ cell differentiation. They showed that proliferation of spermatogonia was reduced in the absence of *hells*, and germ cell differentiation arrested at the midpachytene stage, implicating an essential role for HELLS during male meiosis [63].

As regards to Omy1_b1, there is no direct evidence for a potential role of *myoz1* gene in sex-reversal (see S1 Supplementary Tables, S9 Table). The second gene pointed out for Omy1_b1, *tlx1* (also known as *hox11*), encodes a nuclear transcription factor that belongs to the NK-linked or NK-like (*nkl*) subfamily of homeobox genes. The encoded protein is essential for cell survival during spleen development in fish [64] as in mammals [65, 66], and is also involved in specification of neuronal cell fates during embryogenesis (see S1 Supplementary Tables, S9 Table). At the molecular level, depending on the cellular context and its interaction with transcriptional cofactors, *tlx1* can either activate or repress gene transcription [67]. *tlx1*-dependent regulation of retinoic acid (RA) metabolism is critical for spleen organogenesis [68]. RA is the active metabolite of vitamin A that is required for vertebrate embryogenesis [69, 70]. In mice, the transcription factor steroidogenic factor 1 (*nr5a1* also known as SF1) regulates RA metabolism during germ cell development [71]. *nr5a1* is a transcriptional activator that is essential for sexual differentiation and formation of the primary steroidogenic tissues. Mutations within *nr5a1* underlie different disorders of sexual development, including sex reversal. In patients with disorders of sexual development, a mutant form of *nr5a1* was shown to be defective in activating *tlx1* transcription [72]. In consequence, we can hypothetize that genetic variants in *tlx1* may also directly deregulate RA metabolism, inducing sex-reversal.

Gathering all the information presented here above for the QTL on Omy1, we can conclude that there is a strong statistical and biological evidence of significant involvment in the sex determination and differentiation cascade of *syndig1*, *tlx1*, *hells* and *gbf1*, and, putatively also of *vsx1*, and *myoz1*. Notably, at the only exception of *myoz1*, all these proteins are relatively well conserved in terms of protein identity across various fishes and mammals (Table 5).

### *arfgef3* as the best functionnal candidate for Omy20_a due to its activation of the estrogen/ERα signaling pathway

*arfgef3* gene (named also *big3*) is a member of the *big1*/*sec7p* subfamily of ADP ribosylation factor-GTP exchange factors (ARF-GEFs). The protein is very well conserved across vertebrates (Table 5). *arfgef3* overexpression has been pointed out as one of the important mechanisms causing the activation of the estrogen/ERα signaling pathway in the hormone-related growth of breast cancer cells [73]. As summed up in the introduction, the germ cells seem to play a critical role in sex determination, especially in the feminization of gonads. PGCs may first interpret internal genetic or external environmental cues and directly transform into oogonia; then, the surrounding somatic cells may be induced by the germ cells to differentiate accordingly to provide an appropriate hormonal environment for further gonadal differentiation [11]. In support of this hypothesis, high temperatures were reported to decrease the number of germ cells, which often accompanies female-to-male sex reversal [17]. Guiguen et al. [12] suggested that over-expression of germ cell proliferation or anti-apoptotic germ cell factors in the female gonads could promote ovarian differentiation.

### *khdrbs2*, *dst*, *hmgcll1* and *csmd1* as functional candidates for QTL Omy20_b

#### Implication of khdrbs genes in sex determination and regulation of alternative splicing

In nematodes, *gld*-1, the ortholog of *khdrbs* (KH domain-containing, RNA-binding, signal transduction-associated protein 2), is localized in the germ cell cytoplasm. This gene is indispensable for oogenesis and meiotic prophase progression and it can stimulate sex determination of males in the hermaphrodite germ line [74, 75, 76]. In fruit fly however, *nsr*, the ortholog of *khdrbs* regulates some male fertility genes in the primary [77].

In vertebrates, *khdrbs* (*khdrbs1*, *khdrbs2* and *khdrbs3*) genes encode respectively for the SAM68, SLM1 and SLM2 proteins that have RNA binding and signal transduction activities [78, 79]. In fish, the *khdrbs1* was further duplicated through the teleost-specific whole genome duplication, forming *khdrbs1a* and *khdrbs1b*. In zebrafish, all the *khdrbs* genes were found to be primarily expressed in the brain during early development and at the adult stage. *khdrbs1a* was also found to be expressed in the gonad primordium as well as abundantly expressed in the gonads of adult fish along with *khdrbs1b*, and *khdrbs3* [80]. In our study, eight intronic *khdrbs2* variants were ranked among the 100 best SNPs in DAPC (see S1 Supplementary Tables, S10 Table), and an indel/SNP variant was identified with a p-value < 0.002 in the discovery population (Table 4). In mouse, *khdrbs1* and *khdrbs3* are predominantly expressed in the brain and testis, whether *khdrbs2* is exclusively expressed in the brain [81]. *khdrbs2* knockout mice are fertile fertile whereas *khdrbs1* knockout mice show impaired fertility in males [81]. *khdrbs3* is involved in spermatogenesis via directly binding to RNA [81]. In addition, *khdrbs1 knockout* females displayed a reduction in the number of developing ovarian follicles, alteration of estrous cycles, and impaired fertility (e.g. [82]). It seems that the invertebrate genes *gld-1* and *nsr* that are orthologs of *khdrbs1* and *khdrbs3*, respectively, have similar roles in the regulation of gametogenesis [80].

#### dst and hmgcll1 as novel players in sex determination and differentiation in fish?

*dst* (dystonin) encodes a member of the plakin protein family of adhesion junction plaque proteins. *dst* isoforms are expressed in neural tissue where it regulates the organization and stability of the microtubule network of sensory neurons to allow axonal transport. Low temperatures significantly impact *dst* splicing in killifish, stickleback and zebrafish [83]. Gene ontology annotations related to this gene include calcium ion binding and actin binding (see S1 Supplementary Tables, S9 Table). As far as we know, *dst* has not yet been demonstrated as being involved in sex determination or differentiation. Nevertheless, there is no statistical doubt that this gene is invoved in sex-reversal in rainbow trout. A SNP was highly significant (p-value = 0.0006) in the discovery population (Table 4) with homozygous TT dams exhibiting high ratios of progeny sex-reversal (see S1 Supplementary Tables, S10 Table). In addition, 4 haplotypes within the gene ranked among the 60 best in the full RF analysis (see S1 Supplementary Tables, S8 Table), and 15 intronic SNPs were ranked among the 100 best in DAPC, 10 remaining in the 50 best SNPs (see S1 Supplementary Tables, S10 Table).

*hmgcll1* (3-hydroxymethyl-3-methylglutaryl-CoA lyase-like 1) was first characterized as a lyase activity enzyme localized in extra-mitochondrial region involved in ketogenesis for energy production in nonhepatic animal tissues (see S1 Supplementary Tables, S9 Table). A novel role of *hmgcll1* in cell cycle regulation was recently suggested [84] that could be linked with a putative functional role on sex-reversal. Statistically, there is no doubt about this effect. Two SNPs within the gene were significant in two validation population (Table 3). In the discovery population, an upstream intergenic variant at position 34,345,558 bp was among the 50 best ranked in DAPC, with homozygous TT dams exhibited high ratios of progeny sex-reversal (see S1 Supplementary Tables, S10 Table).

#### csmd1 is associated to gonadal failure in mammals

The *csmd1* (CUB and sushi multiple domains protein 1) gene was one of the best positional candidates in our RF and DAPC analyses with 8 SNPs being among the 50 best ones to discriminate the two groups of extreme phenotypes. This gene encodes for a large (>3,000 amino acids) transmembrane protein. The family of proteins *csmd* (CUB and sushi multiple domains) is one of the three types of the complement system-related proteins involved in the recognition of molecules in innate immune-system and in the central nervous system (reviewed in [85]). *csmd1* is predicted to act upstream of, or within, several processes, including learning or memory, and reproductive structure development. The protein encoded by *csmd1* is highly conserved between vertebrates with 89% amino acid sequence identity between trout and medaka and still 77% between trout and human or mouse (Table 5). In humans and mice, *csmd1* was associated with gonadal failure in both sexes [86]. Significant association between early menopause and rare 5′-deletions of *csmd1* was detected in humans. In mice, knockout females show significant reduction in ovarian quality and breeding success, and *csmd1*-knockout males show increased rates of infertility and severe histological degeneration of the testes. We did not observe any variants predicted with moderate or high impacts on protein expression in this gene. But, as indicated by Lee at al. [86], intronic variants mainly located in introns 1 and 2 likely harbored functional elements that can influence *csmd1* expression in testis.

### *lrrc59* and *caskin2* as the best positional candidates for QTL Omy20_c

As far as we know, no direct effect of *lrrc59* (leucine-rich repeat-containing protein 59) on sex determination is known. This gene enables RNA binding activity and cadherin binding activity (see S1 Supplementary Tables, S9 Table). A key role for *lrrc59* in the organization and regulation of local translation in the endoplasmic reticulum has been suggested [87]. Statistically, it was the best-ranked haplotype on the full RF analysis (see S1 Supplementary Tables, S8 Table), indicating a strong importance of *lrrc59* to predict sex-reversal.

*caskin2* (CASK Interacting Protein 2) encodes a large protein that may be involved in protein-protein interactions (see S1 Supplementary Tables, S9 Table). *caskin2* downregulates genes associated with endothelial cell activation and upregulates genes associated with endothelial cell quiescence [88]. Morpholino knockdown of *caskin2* in zebrafish results in abnormal vascular development characterized by overly branched intersegmental vessels and failure to form the dorsal longitudinal vessels, while *caskin2* knockout mice are viable and fertile [88]. *caskin2* is not well conserved across vertebrates, including only 54% AA identity between rainbow trout and either Medaka or Zebrafish (Table 5). Statistically, two haplotypes including parts of *caskin2* were among the best ranked haplotypes in the full RF analysis (see S1 Supplementary Tables, S8 Table). In addition, 3 SNPs (2 intronic and an upstream intergenic variant) were among the 50 best ranked in DAPC (see S1 Supplementary Tables, S10 Table). The expression of *caskin2* was down-regulated in gonads of *Gobiocypris rarus* of both sexes, when exposed to a 7-day exposure to 17α-methyltestosterone which had a sex-reversal effect [89]. In Coho salmon, androgens were revealed to play a major role in stimulating primary ovarian follicle development and the transition into secondary follicles [90].

### Putative role of *foxl3* in germ cell fate decision

Intriguingly, *foxl3*, a co-paralog gene of *foxl2*, is located on Omy20 spanning from 22,942,515 to 22,948,650 bp, upstream the QTL regions we studied on the same chromosome. As *foxl3* is expressed in the gonads of teleosts, we checked if any variant within *foxl3* could be associated to sex-reversal in our study, and could not find any. However, we can speculate that *foxl3* is involved in germ cell fate decision in rainbow trout. Indeed, *foxl3* was already suggested to be involved in the onset of oocyte meiosis, and/or in the regulation of male specific genes during late testis development and testis maturation in salmonids [91, 92, 93]. It was also demonstrated in medaka that *foxl3* expression is only maintained in female germ cells, and that functional sperm is produced in the ovary of a *foxl3* loss-of-function mutant [94]. In addition, *foxl3* was shown to initiate oogenesis in medaka through two genetically independent pathways, meiosis and follicular development, and that a third pathway indepentdent of *foxl3* may also be hypothetized [95]. Last, *foxl3* expression levels were strongly increased in gonads during the natural female-to-male sex reversal in the rice field eel (*Monopterus albus*). *foxl3* and miR-9 seemed to be involved in physiological processes that promote oocyte degeneration in the ovotestis and stimulating spermatogenesis in spermatogonia [96]. Therefore, we can hypothetize that expression of *foxl3* may be regulated by some other genes, presumably located in the QTL regions identified on Omy20 in our study, similarly to the long-distance cis-regulation observed for *sox9* expression [57]. Further investigation should try to validate such speculations, as well as the causative variants and precise functional roles of the identified genes in our QTL regions.

## Conclusions

While considerable scientific effort has been made over the last fifty years to unravel mechanisms for sex-determination and differentiation, the nature of genetic variations that regulate these mechanisms in fish is still not well-known. We confirmed on several French commercial rainbow trout populations, the importance of some genomic regions from chromosomes Omy1, Omy12 and Omy20 as minor sex-determing genes involved in maleness in the absence of the master sex-determing gene *sdY* in rainbow trout. Together, our results are consistent with a model in which spontaneous female-to-male sex-reversal in rainbow trout is associated with genetic factors able to reduce germ cell proliferation and arrest oogenesis. The main candidate genes that we suggested based on positional and functional information are *syndig1*, *tlx1*, *hells* and *gbf1* on Omy1, and *arfgef3*, *khdrbs2*, *dst*, *hmgcll1*, *csmd1*, *lrrc59* and *caskin2* on Omy20. Most of these genes were previously indicated as playing roles in developmental pathways in vertebrates.

From a fish farmer perspective, further investigation is still needed before any operationnal use of the genotypes at minor sex-modifier genes either to eradicate spontaneous sex-reversal in all-female production stocks or to produce progeny from free-hormone neomales in rainbow trout. Indeed, the high heritability of maleness [27] suggests that the use of spontaneously masculinised individuals as progenitors of all-female stocks would increase the frequency of undesirable masculinised progeny. Improve knowledge about the joint effects of genetic basis and environmental factors determining spontaneous maleness in all-female stocks is then needed to search for a trade-off combining together genetic and environmental control of gonad masculinisation according to the destination of the fish (broodstock vs all-female production stock). High temperature during development masculinizes many fish species [97]. In rainbow trout, high rearing temperature during the early phase of fry development after hatching was shown to increase the frequency of sex reversal in XX females to a greater or lesser extent depending on the genetic background of the trout lines [98, 99]. Further investigation should therefore focus on the interactions between genetics and rearing temperatures to determine the key combination of factors to produce all-female progeny from free-hormone neomales in rainbow trout.

## Supplementary Information

**S1 Appendix (S1-Appendix.doc)**

**Positional candidate genes on Omy12 and Omy20 without known evidence of a functional role linked to sex-reversal**

**S1 Supplementary Tables (S1-Supplementary Tables.xls). This excel file includes 10 supplementary tables whose titles are listed below.**

**S1 Table Number of progeny (female, intersex and male) and proportion of neomales per sequenced dam**

**S2 Table Positions on the Swanson reference genome and flanking sequences of the 192 SNPs used in validation**

**S3 Table List of the 192 SNPs detected in 6 validation populations (named A to F) with their positions on Swanson (SW) and Arlee (AR) reference genomes**

**S.4 Table List of most significative SNPs found in Omy1, Omy12 and Omy20 QTL**

**S5 Table Mean, Minimum and Maximum values of the 30 haplotypes with the highest Gini index over 100 runs of Random Forests for Omy1 QTL**

**S6 Table Mean, Minimum and Maximum values of the 10 haplotypes with the highest Gini index over 100 runs of Random Forests for Omy12 QTL**

**S7 Table Mean, Minimum and Maximum values of the 30 haplotypes with the highest Gini index over 100 runs of Random Forests for the 3 Omy 20 QTL (Z3, Z4, Z5)**

**S8 Table Mean values of the 60 haplotypes with the highest Gini index over 100 runs of Random Forests for the QTL regions of Omy1 (Z1), Omy12 (Z2) and Omy20 (Z3 to Z5)**

**S9 Table List of potential candidate genes for QTL on Omy1, Omy12 and Omy20 with their descriptions on GeneCards, UniProtKB/Swiss-Prot and NCBI databases.**

**S10 Table Location, annotation, alleles and genotypes for sequenced dams of the 100 most important SNPs in DAPC analysis. Genotypes in blue are considered as male genotypes, genotypes in orange as female genotypes and genotypes in light orange are heterozygous.**

## Acknowledgements

The European Maritime and Fisheries Fund and the French National Government supported this work (R FEA470016FA1000008). The authors are grateful to the companies “Les Fils de Charles Murgat”, “Aqualande” “Bretagne Truite”, “Font Rome” and “Viviers de Sarrance” for fish samples and data collection.

## Author contributions

AD: Data curation, Formal analysis, Preparation of tables and figures, Writing—review and editing. CF: Data curation, Formal analysis, Preparation of figures, Writing—review and editing. AB: Data collection, Writing—review and editing. CP: Data collection. YG: Data collection, Interpretation, Writing—review and editing. EQ: Conceptualization, Data collection, Funding acquisition, Project administration, Writing— review and editing. FP: Conceptualization, Methodology, Supervision, Statistical analysis, Interpretation, Project administration, Writing—original draft, review, and editing. All authors read and approved the final version of the manuscript.

## Data availability

Raw sequence data that were used in this study are deposited in the ENA (Project PRJEB75960 in https://www.ebi.ac.uk/ena/browser/view/PRJEB75960).

## Declarations

### Animal and human rights statement

We declare that all applicable international, national, and/or institutional guidelines for sampling, care, and experimental use of organisms for the study have been followed and all necessary approvals have been obtained. As part of the standard breeding practices for a commercial breeding program, the handling of fish was not subject to oversight by an institutional ethic committee. Animals were treated in compliance with the European Communities Council Directive (9858/CEC) for the care and use of farm animals.

